# LAWS: Local Alignment for Water Sites — tracking ordered water in simulations

**DOI:** 10.1101/2022.05.12.491708

**Authors:** Eugene Klyshko, Justin Sung-Ho Kim, Sarah Rauscher

## Abstract

Protein solvation plays a crucial role in protein function; accurate modeling of protein-water interactions is important for understanding the molecular basis of protein function. In addition to protein structure, x-ray crystallography provides positions of crystallographic water sites (CWS) coordinated by the protein, which can serve as a strong benchmark for modeling accuracy. Such comparison requires special methodological considerations that take into account the dynamic nature of proteins. However, existing methods for analyzing CWS in MD simulations rely on global alignment of the protein onto the crystal structure, which introduces substantial errors in the case of significant structural deviations. Here, we propose a method called local alignment for water sites (LAWS), which is based on multilateration — an algorithm widely used in GPS tracking. LAWS considers the contacts formed by CWS and protein atoms in the crystal structure and uses these interaction distances to track the CWS in a simulation. LAWS provides a framework to quantify how perturbations of the local protein environment affect the preservation of crystallographic waters in simulations. We applied our method to a 1 *µ*s simulation of a protein crystal and demonstrate that 76% of CWS are preserved. Compared to existing methods, LAWS identifies more high-confidence (low B-factor) CWS, which are characterized by more prominent water density peaks and a less-perturbed protein environment.

## Introduction

Water is an essential component of biomolecular systems; it affects the structure and stability of biological machinery through the hydrophobic effect, hydrogen bonding, and polar interactions.^1^ Protein-water interactions are important driving forces in dynamic processes, such as protein folding, self-assembly, and binding. ^2,3^ In addition, solvation is essential for protein function, since water networks have been shown to play a crucial role in the motion of protein domains.^1,4–6^ All-atom molecular dynamics (MD) simulations can provide information on the molecular basis for protein function, but require high-resolution structural information.^1,7^

X-ray crystallography is the most commonly used experimental technique to obtain high-resolution protein structures.^8^ Crystal structures often contain crystallographic water sites (CWS), which are locations of high water density in the lattice. These CWS represent ordered water molecules, which are stabilized by non-covalent interactions with the protein, while bulk (unordered) water molecules comprise the remaining space between the protein chains. This crystallographic information provides a direct benchmark to assess the accuracy of modeling protein-water interactions.^9,10^ MD simulations can probe the dynamics of the protein crystals because periodic boundary conditions mimic the periodic nature of crystal lattices.^9,11,12^ However, since x-ray crystallography provides an ensemble-averaged conformation of the protein and CWS, a direct comparison between time-resolved MD trajectories and experiment requires special methodological considerations.

The analysis of CWS in MD simulations has been carried out using a variety of methods that can be broadly classified into two categories — density-based and coordinate-based methods (Fig. 1). In density-based methods,^13–20^ the time-averaged atomic density of the solvent molecules is computed, where high-density peaks correspond to CWS and are compared to the experimental CWS positions (Fig. 1A). In contrast, the coordinate-based methods^13,17,21–23^ analyze the explicit positions of the water molecules in each MD frame in the vicinity of CWS (Fig. 1B and 1D). To identify if CWS are preserved in simulations, their locations are probed for the occupancy of water molecules. We refer to these locations in the MD trajectory as water sites (WS); they can be tracked in various ways depending on whether the protein conformation is globally or locally aligned to the crystal structure. When using global alignment (Fig. 1A and 1B), the entire protein is superimposed onto the crystal structure and the experimental positions of the CWS represent the WS in a frame. In the local alignment approach, WS are defined using only a local region of the protein in the vicinity of the experimental CWS (Fig. 1D).

**Figure 1:**
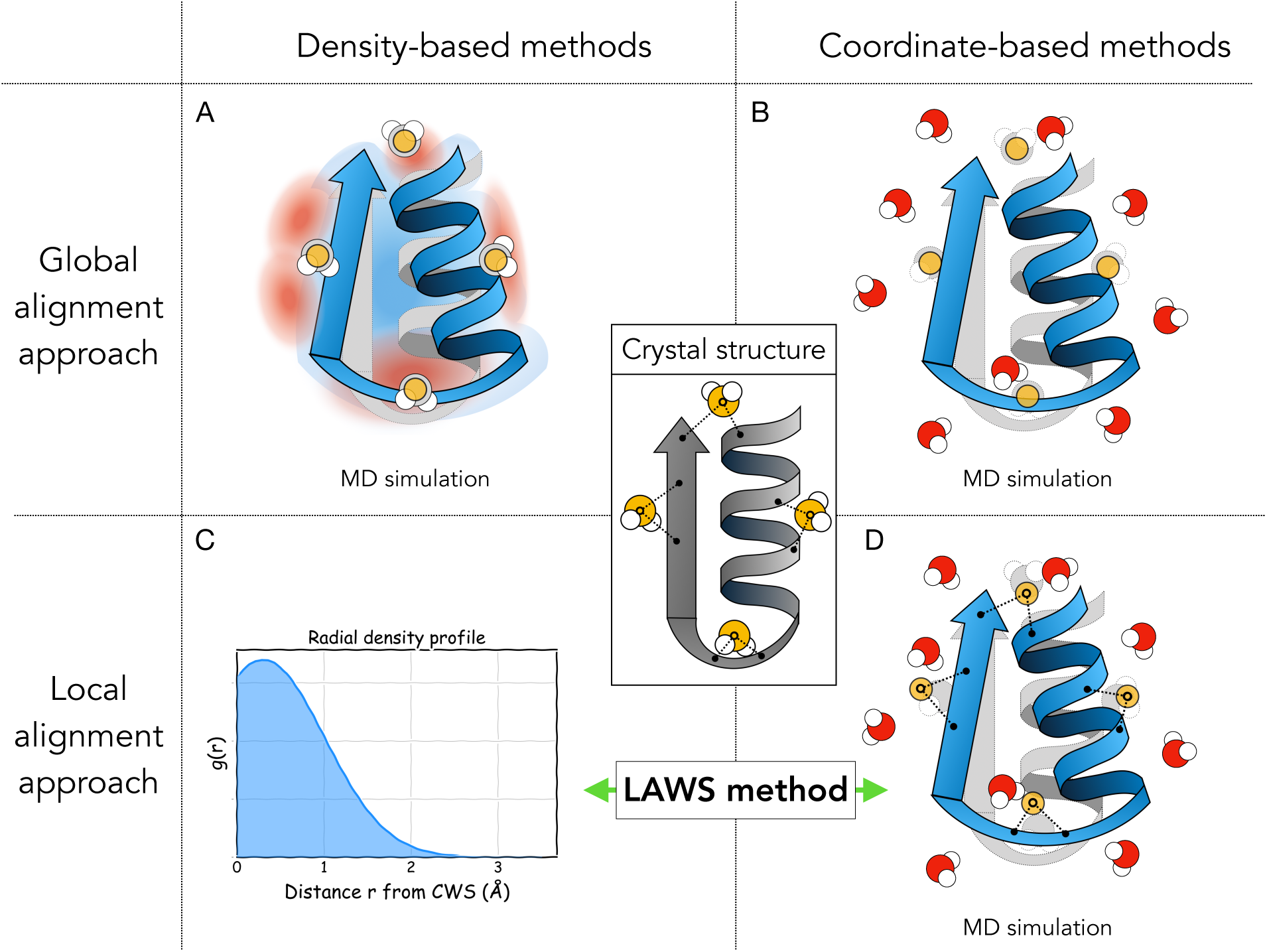
A schematic illustration of existing methods for the analysis of crystallographic water in MD simulations. These methods are based on either average MD density maps or explicit MD coordinates, and can rely on global (**A**,**B**) or local (**C**,**D**) alignment of the protein. The crystal structure (central panel) provides the experimental positions of CWS (yellow spheres). Density-based methods (**A**) analyze an average atomic density of water in an MD simulation and then compare CWS to the peaks in the density (red). Coordinate-based methods (**B, D**) analyze explicit positions of water molecules in every frame and measure the occupancy of each CWS, which can be done using either global (**B**) or local (**D**) alignment of the protein structure to the crystal structure. Global alignment approaches (**A, B**) superimpose the configuration of the protein from an MD simulation (blue) onto the static crystal structure (grey), which can result in clashes between CWS and protein atoms. Local alignment approaches (**D**) address this issue by only considering local protein regions in the vicinity of the CWS. The method proposed here — Local Alignment for Water Sites (LAWS) — is a coordinate-based algorithm relying on a local alignment approach, which additionally provides density-type data, including the local radial density profile (**C**) or density maps (Fig. S3). Therefore, LAWS can be classified as both a coordinate-based and a density-based method (**C, D**).

In the analysis of CWS, a quantity of interest is the proportion of CWS preserved in a simulation, referred to as *recall statistics*.^13–15,18,21,24^ It has been demonstrated that density-based and coordinate-based methods produce comparable recall of CWS when global alignment is employed. ^13^ However, the global alignment approach has one notable limitation; it is only suitable for analysis of short (tens of ns) or restrained MD simulations in which the protein does not deviate significantly from the crystal structure. ^13–15,18,21^ Even in a crystal environment, proteins are flexible and can undergo conformational changes at sub-*µs* timescales.^10,25,26^ For example, Wall et al. showed that 93% of all CWS were preserved within 1.4 Å of their experimental positions in a simulation with position restraints, while only 46% of CWS were preserved when position restraints were removed, suggesting that the deviation of a protein from its crystal structure may contribute to the loss of CWS in simulations.^14^ Significant changes in the local protein structure near a CWS will perturb protein-water interactions, resulting in shifts of the corresponding CWS in each frame of the simulation (Fig. 1D). Such effects motivate the local alignment approach.

To analyze CWS in simulations, Henchman and McCammon developed a local alignment approach using backbone atoms.^17^ However, side chain atoms also contribute to the coordination of crystallographic waters. Caldararu et al. proposed a method that includes local side-chain atoms in the alignment to track CWS. This method relies on clustering coordinates of the water molecules located near CWS.^13^ However, clustering algorithms have their limitations as they are data-driven and can be sensitive to the choice of parameters.^27^ A natural way to analyze CWS in MD simulations is by using a measure of density, as this data is probed in x-ray crystallography experiments.

Here, we propose a method to analyze CWS called local alignment for water sites (LAWS). LAWS is based on a widely used algorithm in GPS navigation called multilateration^28^ to track CWS relative to nearby protein atoms in an MD trajectory. LAWS has several advantages: (i) it reduces alignment errors compared to traditional global alignment approaches, (ii) it quantifies the local structural perturbations in the vicinity of CWS, and (iii) it provides local density information for each CWS. We apply the LAWS method to a 1 *µ*s MD simulation of a protein crystal. LAWS is found to characterize CWS with a higher density of water compared to the global alignment approach. We also explore how various properties of the CWS, such as the experimental uncertainty (B-factors) and the location relative to the protein, affect the recall of CWS in the simulation.

## Methods

### The LAWS algorithm

In a crystal structure, each CWS can be characterized by a set of contacts with nearby protein atoms. When similar interactions are maintained in a simulation, the position of the crystal water would be preserved over time. Therefore, in the LAWS algorithm, we track water sites using these interaction distances and compute the density of water molecules around each water site (yellow spheres in Fig. 2A).

**Figure 2:**
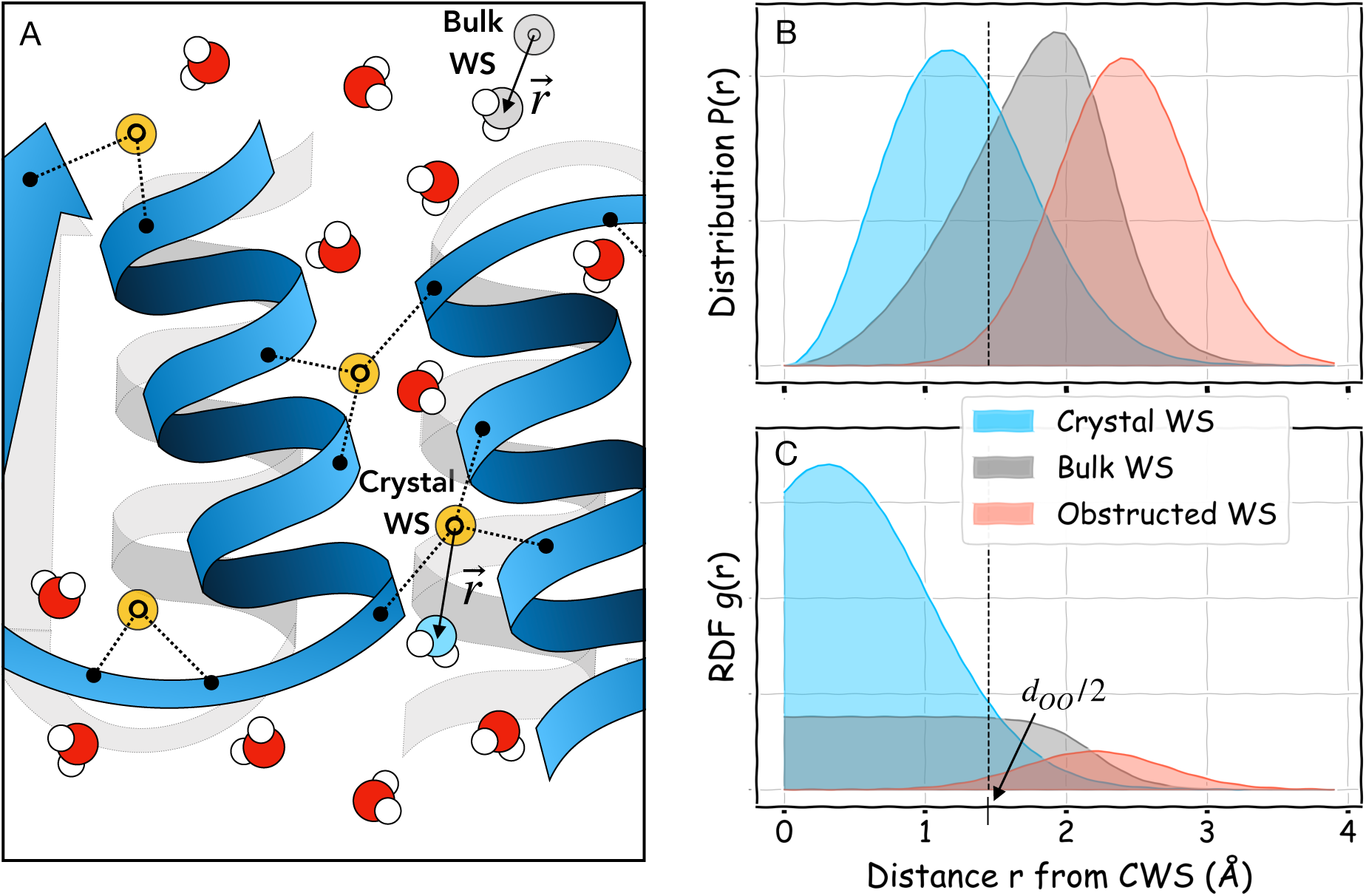
The LAWS pipeline. (**A**) A cartoon representation of one MD frame containing positions of protein and water molecules, overlapping with a crystal structure in grey. Yellow points represent water sites which are tracked by the LAWS algorithm based on the distances to nearby protein atoms. Note that contacts can be formed by multiple protein chains. To check if water sites are crystallographic and preserved in MD, we compute the distance *r* to the closest water molecule at each frame. Statistically, these sites are more likely to be occupied than bulk water sites shown in grey. (**B**) The resulting distance distributions *P* (*r*) for a preserved CWS, bulk WS, and obstructed WS. (**C**) The resulting radial distribution functions *g*(*r*) = *P* (*r*)/*r*^2^. A radial density around bulk water sites must be uniform from zero to *d*_*OO*_/2 (shown as a dashed line), where *d*_*OO*_ is the average distance between nearest neighbour oxygen atoms in bulk water.

#### Tracking water sites in a simulation using LAWS

Let *x, y* and *z* be the unknown coordinates of a water site in a simulation frame, and *A*_*i*_ are the *n* nearby protein atoms with coordinates *x*_*i*_, *y*_*i*_, *z*_*i*_ and indices *i* = 1, …, *n*. Using the coordinates in the crystal structure, the distance between an atom *A*_*i*_ and the water site, 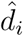, is initially determined (dashed lines in the central panel of Fig. 1). The set of fixed distances 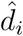 to the nearby protein atoms can then be used to find the unknown coordinates of the WS while the protein atom positions *x*_*i*_, *y*_*i*_, *z*_*i*_ change at every frame. In other words, for every frame we are aiming to find the position *x, y, z* of the WS that satisfies the *n* distance equations:

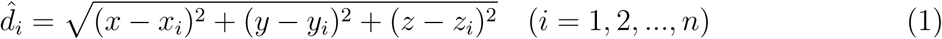

This problem is formulated in the literature as multilateration,^28^ which is commonly encountered in navigation and surveillance. For example, a GPS device calculates distances 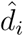 by measuring the times required for a signal to travel from a set of satellites. Since the position of each satellite *x*_*i*_, *y*_*i*_, *z*_*i*_ is known at any given time, the coordinates of the GPS device *x, y, z* can be determined by solving a system of equations (1). By analogy, the protein atoms act as the satellites and the GPS-device represents a CWS with a location that needs to be determined.

The system of non-linear equations (1) has three unknowns and *n* equations. In theory, the exact solution of the equation can be found uniquely if *n* = 4 positions and distances are provided — analogous to identifying the point of intersection of four spheres with known radii and positions of the centres in 3D space. However, an ambiguity of the solution is possible when any two sphere centres and the unknown point are collinear. In that case, more spheres are required for a unique solution. On the other hand, Eq. (1) is not guaranteed to have a solution as the spheres may not intersect in every case. Thus, due to the stochastic nature of protein motions and the fact that it is not possible to find an exact solution in every instance, an *optimal* solution is required instead.

We define the LAWS error as the weighted sum of squares:

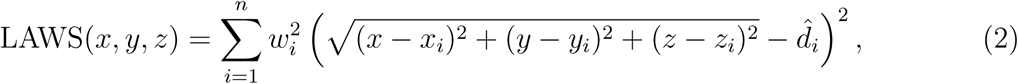

where *w*_*i*_ are the weights of each atom *A*_*i*_ in defining the water site, such that 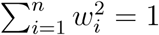. This function is then minimized to find the optimum position *x*^*^, *y*^*^, *z*^*^ of the water site in a given simulation frame:

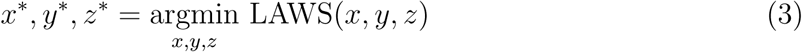

This procedure is the weighted least-squares problem for which we used a Python implementation of the Levenberg-Marquardt algorithm. ^29^

The motivation for using weights comes from the susceptibility of the least-squares method to outliers. Protein atoms closer to the CWS in the crystal have a higher contribution to the interaction energy than atoms further away. Hence, they have greater weights in the LAWS error (2), which increases the robustness of the algorithm. Importantly, the value of the LAWS error at the optimum *x*^*^, *y*^*^, *z*^*^ shows a quantitative estimate of how much the local protein region is perturbed relative to the crystal structure. The exact solution will have a LAWS error of zero. Therefore, the LAWS error is a measure of the displacement of the WS from its ideal position in the crystal structure.

#### Parameters

When applying the LAWS algorithm to an MD trajectory, the number of protein atoms, *n*, coordinating the CWS and the weights *w*_*i*_ must be determined. These parameters are computed only once based on the crystal structure.

There is a trade-off between computational efficiency and robustness of the numerical algorithm that depends on the number of protein atoms *n* used to define each CWS. A larger *n* increases the computational cost of solving Eq. (3), while a smaller *n* makes the algorithm less robust to outliers. The value of *n* depends on the cutoff distance defining a contact. We use a cutoff distance of 4.5 Å and set a maximum of *n* = 10. If there are fewer than four protein atoms in contact with the CWS, we increase the cutoff distance until we find at least *n* = 4. Therefore, *n* varies between 4 and 10 depending on the CWS. Importantly, we take into consideration protein atoms from multiple symmetrically related molecules if the CWS is located at the interface between molecules in the unit cell (Fig. 2A). We also tested the LAWS algorithm with the same fixed number of heavy atoms *n* for all CWS in a range from 5 to 8 and the results were not significantly affected by this parameter choice (data not shown). Once *n* protein atoms are determined, the distances 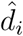 can be computed and used in Eq. (2).

The choice of weights *w*_*i*_ in Eq. (2) is motivated as follows: the protein atoms located closer to the WS contribute more to its coordination. The protein-water interaction energy depends on the distance, *r*, between atoms as ∝ 1/*r*^*α*^, where *α* ≥ 1 for various non-covalent interactions. We chose the weights *w*_*i*_ to be proportional to the interaction energy and hence inversely proportional to the distance between the CWS and a protein atom in the crystal structure (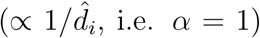). We tested other values of *α* (*α* = 2, …, 6) and this did not affect the results significantly. The normalized weights are therefore given by:

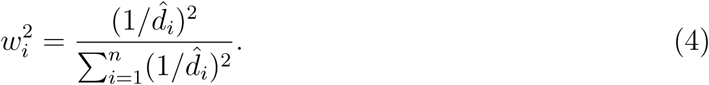

### Computing density profiles of water sites

To estimate the occupancy of any WS, we compute the offset vector 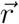 and the distance *r* from a WS to the oxygen of the nearest water molecule at each frame. The distance distribution *P*(*r*) (averaged over all symmetrically-related copies) then provides a measure of the occupancy of the WS in the trajectory, as it shows the fraction of nearest neighbour water molecules found at the spherical shell with radius *r* from the WS (Fig. 2B). Since the probability to find water at a spherical shell with radius *r* is proportional to the area of a sphere, *P*(*r*) ∝ 4*πr*^2^, we normalize distribution *P*(*r*) by *r*^2^ to define a radial distribution function (RDF). Then, *g*(*r*) = *P*(*r*)/*r*^2^ provides a density profile as a function of the distance *r* from the WS (Fig. 2C). Additionally, offset vectors 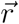 around each WS can be used to generate local 3D density maps with, for example, GROma*ρ*s software.^30^ The details of this approach are presented in Supporting Information (SI Section 4).

#### Bulk water sites as reference

A CWS in a simulation is expected to have on average a higher occupancy than a WS in a bulk water region, which can be treated as a reference for comparison. ^31^ Bulk WS are defined as positions within the unit cell where water molecules are not significantly influenced by the interaction with the protein. Water was experimentally observed to have bulk-like properties (in terms of the lifetime of hydrogen bonds) at least 6 Å from the protein surface. ^32^ Based on this observation, we checked if randomly sampled positions in a unit cell located at least 6 Å from the protein would display bulk water properties (grey spheres in Fig. 2A). We ran a 500 ns simulation of a box of water to model ideal bulk water behaviour (SI Section 1). Analyzing the *P*(*r*) distribution for randomly sampled locations in the box, we found that the bulk water *P*(*r*) was statistically identical to the *P*(*r*) sampled at 6 Å from the protein in our MD simulation of a unit cell (Fig. S1). Therefore, water located > 6 Å from the protein surface exhibits bulk behaviour, and hence the *P*(*r*) from the unit cell simulation can be used as the reference bulk distribution.

The radial density around a bulk WS should be uniform from zero to *d*_*OO*_/2 (Fig. 2C), where *d*_*OO*_ is the average inter-oxygen distance between nearest neighbour water molecules in liquid water. Hence, it is equally likely to find the nearest neighbour water molecule at any distance *r* ≤ *d*_*OO*_/2 from a bulk WS. When *r > d*_*OO*_/2, the probability to observe a nearest neighbour water becomes negligible as *r* approaches *d*_*OO*_. The experimental estimate of *d*_*OO*_ was reported to be in the range of 2.7-3.0 Å.^33,34^ We estimated a value of *d*_*OO*_ = 2.8 Å using the *g*(*r*) from our simulations, consistent with previous MD studies. ^14,15^

#### Crystal, bulk-like, and obstructed water sites in a simulation

Given the RDF *g*(*r*) for each WS, we can classify it into one of three groups: crystal, bulk-like, or obstructed. The radial density profile for a crystal WS is expected to have a peak, the radial density for a bulk-like WS should be uniform, while the radial density of an obstructed WS is shifted to the right (Fig. 2C). Obstructed WS occur when the protein or another cosolute occupies this space, obstructing access of water molecules. The metric we used to compare the RDF to the bulk reference is the integral 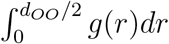 of the RDF in the interval from 0 to *d* /2. First, we computed this integral for randomly sampled bulk water sites to estimate a range (*I*_*MIN*_, *I*_*MAX*_) for comparison. Then, if the RDF integral for a WS is greater than *I*_*MAX*_, we consider this CWS to be preserved in the simulation (Fig. 2C, blue). If the WS has an RDF integral in the range (*I*_*MIN*_, *I*_*MAX*_), then the WS is not preserved in the simulation and is classified as a bulk-like WS (Fig. 2C, grey). Lastly, an RDF integral below *I*_*MIN*_ indicates that the WS is obstructed in the simulation and also not preserved (Fig. 2C, red).

### Global Alignment vs LAWS comparison

We compared global alignment and LAWS as two approaches for tracking positions of CWS in a simulation. In global alignment, for each frame of the trajectory, the protein structure was aligned to the crystal structure by minimizing RMSD using the MDAnalysis Python library.^35^ In this case, positions of aligned crystallographic water oxygen atoms defined water sites. Once the WS are defined at each simulation frame (using either global alignment or LAWS), we can compute the occupancy of these sites using the RDF approach described above. Globally aligned CWS positions are fixed relative to each other, whereas in LAWS they can move relative to each other due to changes in the protein structure.

### MD Simulation Details

#### Model building

The simulation system was constructed from the 95-residue long second PDZ domain of the ligand of Numb protein X 2 (LNX2^PDZ2^) which was obtained from PDB entry 5E11^25^ with a resolution of 1.80 Å (Fig. 3). We chose this structure as an example of a room temperature protein crystal structure. Alternate conformations with the highest occupancy were chosen in model building. We used the CHARMM-GUI web-server to add atoms that were missing in the crystal structure.^36^ The positions of all hydrogen atoms in the original PDB file were ignored. Instead, hydrogen atoms were added to the initial structure using *pdb2gmx* in the GROMACS software package.^37^ Next, CHARMM-GUI^36^ was used to reconstruct a triclinic unit cell (*C*121 space group, with parameters *a* = 65.30 Å, *b* = 39.45 Å, *c* = 39.01 Å and *α* = *γ* = 90^°^, *β* = 117.54^°^) with four symmetrically-related proteins (Fig. 3). Since the conditions (salt concentration, pH, etc) of the protein crystal are difficult to determine, these parameters were chosen to match the conditions of the crystallization buffer as closely as possible. Sodium chloride (NaCl) was added to neutralize the system and then to mimic the 35 mM concentration of NaH_2_PO_4_ found in the buffer. To mimic the pH of the crystallization buffer (pH 4.5), the residues were matched to their protonation states at this pH: N- and C-termini were kept charged and all of the histidine residues were protonated.^25^ All of the crystallographic water oxygen atoms (with the highest occupancy in case of alternate conformations) were included in the construction of the unit cell. Following the method of Cerutti and Case,^11^ additional water molecules were added to solvate the system and preserve the experimental volume of the unit cell in the NPT ensemble using the GROMACS utility *solvate*.^37^ The simulation system contained a total of 9650 atoms, with 6892 protein atoms, 3750 water atoms, as well as 6 sodium ions, and 2 chloride ions.

**Figure 3:**
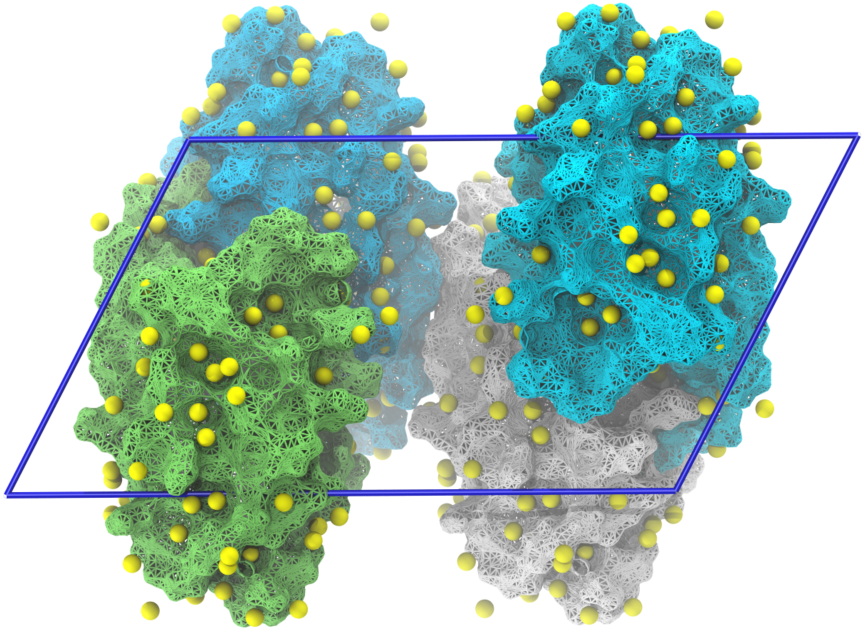
The simulation system. A single unit cell of the LNX2^*P DZ*2^ crystal with the *ac*-plane shown. There are 94 CWS (yellow spheres) around each of four symmetrically related protein chains (coloured individually and shown in a surface representation), with a total of 376 CWS.

#### Simulation

Simulations were conducted using GROMACS 2019.1.^37^ The CHARMM36m force field^38^ combined with the CHARMM-modified TIP3P water model^39^ were used for the study. The time step of the simulation was 2 fs. The LINCS algorithm was used to constrain covalent bonds with hydrogen atoms. ^40^ Short-range electrostatics and Lennard-Jones interactions were computed with a cutoff of 9.5 Å. Long-range electrostatics were computed using particle-mesh Ewald summation with a grid spacing of 1.2 Å with a fourth order interpolation.^41^ A compressibility of 2.5 × 10^−5^ bar^−1^ was used to mimic the compressibility of a protein crystal.^42^ The temperature of the simulation was kept constant at 298 K using the velocity rescaling thermostat^43^ to match the experimental conditions. The pressure of the system was kept constant at 1 bar. Following energy minimization of the system using the steepest descent algorithm, 10 ns of position restrained simulation was performed, followed by equilibration in the NVT ensemble for 100 ns. Two types of NPT equilibration simulations were performed in succession: (i) 10 ns of isotropic Berendsen pressure coupling to quickly reach a pressure of 1 bar and (ii) 100 ns of simulation using isotropic Parrinello-Rahman pressure coupling.^44,45^ The simulation was extended for an additional 1 *µs*, which was used for the analysis (100,000 frames with a 10 ps stride). The protein reached a heavy-atom root-mean-squared deviation (RMSD) of 1.75 Å relative to the crystal structure (Fig. S2).

#### Crystallographic water sites

In the crystal structure (PDB 5E11), there are a total of 94 CWS. Each CWS is represented by the position of an oxygen atom with the highest occupancy. Thus, we tracked 94 WS associated with each of the four individual PDZ domains in the unit cell, such that that there are four symmetric copies of each CWS, providing a total of 94 × 4 = 376 WS for the entire unit cell (Fig. 3). A CWS was classified as intra-chain (*n* = 38) if its coordination was limited to a single protein chain, whereas it was classified as inter-chain (*n* = 56) if it was coordinated by multiple distinct chains of the unit cell.

## Results and Discussion

### Recall statistics of CWS

We applied the LAWS pipeline (described in Methods and summarized in SI Section 2) to the 1 *µs* simulation of the PDZ domain unit cell (Fig. 3), which contains four symmetrically related copies of the protein. Additionally, we performed a similar analysis using the global alignment approach to define the WS in each frame: by superimposing the crystal structure onto the simulation structure, the aligned CWS represent the WS positions (Fig. 4A). After we tracked the WS in the simulation (using either LAWS or global alignment) and computed the RDF for each WS, we classified each WS into three groups according its RDF: crystal, bulk-like, and obstructed (Fig. 4B).

**Figure 4:**
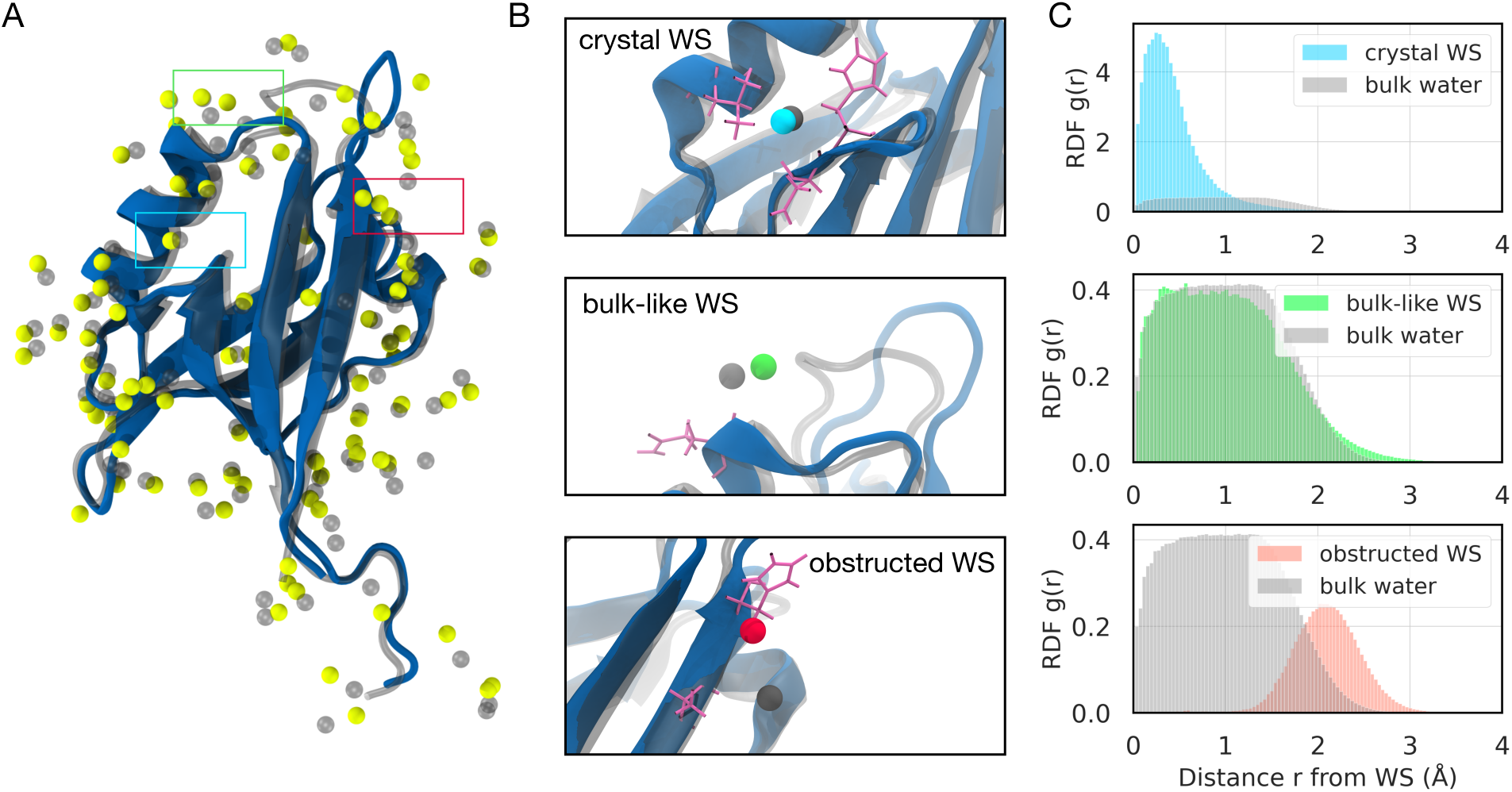
The LAWS method classifies water sites into crystal, bulk-like and obstructed. **(A)** A representative structure of the PDZ domain taken from the simulation (blue) is superimposed onto the crystal structure (grey). Water sites obtained by LAWS (yellow) are compared to the water sites obtained by global alignment (grey spheres). Some of the water sites are coordinated by interactions with multiple protein chains in the unit cell, however only a single protein is shown for clarity. **(B)** The local structure surrounding a representative crystal WS (blue), bulk-like WS (green), and obstructed WS (red). In each panel, the positions of globally aligned WS (grey spheres) and the coordinating side chains (purple) are shown. **(C)** Radial distribution functions for each representative water site in (B) compared to the control bulk water distribution are shown.

Crystal WS are characterized by a high radial density of water in the range from 0 to 1.4 Å when compared to the control bulk distribution (Fig. 4C, top). Bulk-like WS have an RDF which looks like that of the control bulk water, corresponding to the regions of average density (Fig. 4C, center). Obstructed WS are located close to the protein (within the Van der Waals radii of protein atoms), preventing water molecules from occupying these WS regions. This obstruction results in the RDF shifting to the right (Fig. 4C, bottom).

We computed the number of CWS classified as crystal, bulk-like, and obstructed in the simulation using both LAWS and global alignment (Table 1). Only the WS classified as crystal WS are considered to be preserved in the simulation, whereas bulk-like and obstructed WS are not preserved. Using LAWS to analyze the simulation, we find that 76% of the CWS are preserved. The results with the global alignment approach shows a smaller CWS recall: 68% are identified in the simulation as preserved.

**Table 1:** The number of water sites (with percentage) classified as preserved, bulk-like and obstructed using LAWS and global alignment methods

| Method | Preserved CWS | Bulk-like WS | Obstructed WS |
| --- | --- | --- | --- |
| LAWS | 71 (76%) | 10 (11%) | 13 (14%) |
| Global Alignment | 64 (68%) | 8 (9%) | 22 (23%) |

To test the robustness of our method, namely, estimating occupancy with radial density *g*(*r*), we analyzed the 3D density peaks computed from the offset vectors 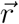 (Fig. S3). The CWS recall obtained using the radial density does not differ significantly from the results obtained using 3D density maps (Table S1). Irrespective of the method used to compute the CWS occupancy, LAWS shows more CWS to be preserved in the simulation when compared to global alignment.

### LAWS detects CWS with higher density compared to global alignment

To compare the properties of the CWS identified with LAWS with those identified by the global alignment procedure, we compute the distance distribution, *P*(*r*), combined for all preserved CWS (Fig. 5A), and estimate the RDF *g*(*r*) (Fig. 5B). An apparent shift of the *P*(*r*) distribution to smaller *r* is evident for the preserved crystal WS identified by LAWS compared to global alignment (Fig. 5A). This shift leads to a peak in the radial density that is significantly higher (Fig. 5B). The analysis of 3D density maps demonstrates similar results; CWS defined by LAWS have higher 3D density peaks than those defined by global alignment (Fig. S4). Therefore, water is more localized around the CWS tracked by the LAWS algorithm than by global alignment.

**Figure 5:**
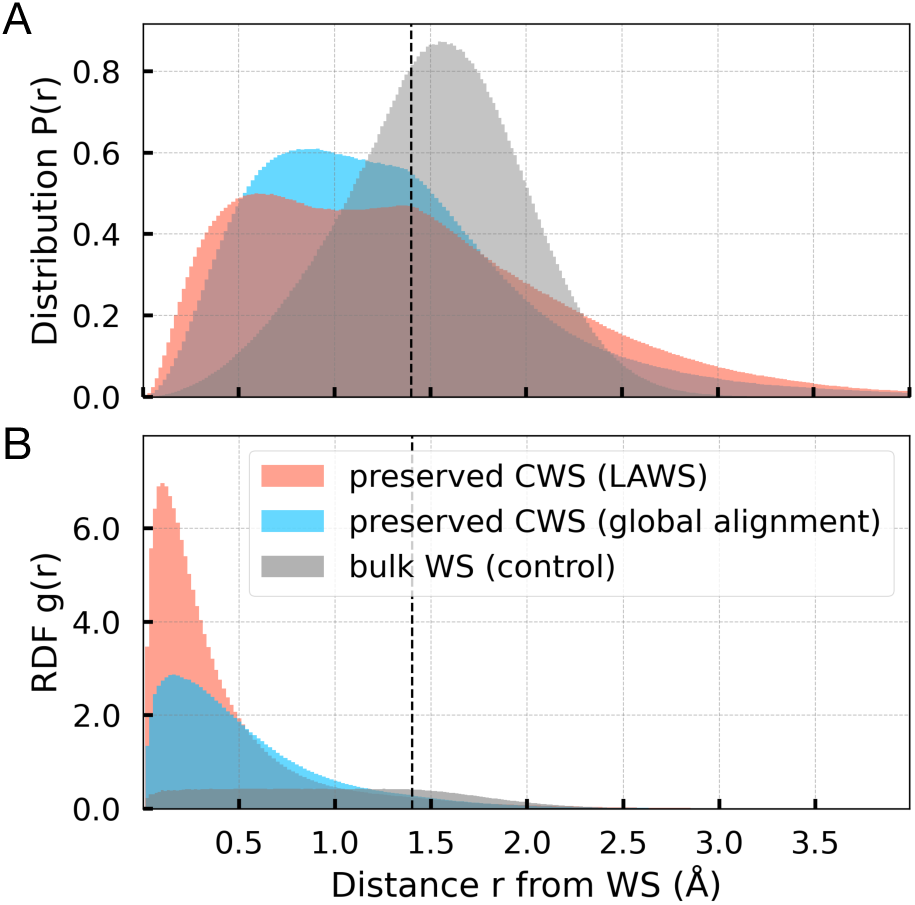
CWS detected by LAWS exhibit higher water density compared to global alignment. **(A)** Distribution of distances, *P* (*r*), between the nearest neighbour water molecule and (i) CWS tracked with LAWS (orange), (ii) CWS tracked with global alignment (blue) and (iii) bulk WS (grey). **(B)** Corresponding RDF, *g*(*r*), with the same colour scheme as **(A)**. For instance, it is ∼ 7× more likely relative to bulk to find a water molecule within 0.2 Å from the CWS, as found by global alignment. In contrast, it is ∼ 17× more likely, as found by LAWS. The likelihood of finding a water molecule within 0.2 Å from the CWS relative to bulk is increased by ∼ 2.4× by using LAWS versus global alignment.

### LAWS quantifies perturbation to the local protein structure

To determine how the deviation of the protein from the crystal structure contributes to the loss of crystal water, we analyzed the LAWS error (Eq. 2) for each group of water sites. The LAWS error represents a quantitative measure of the perturbation to the protein region coordinating the CWS in a given MD frame. A LAWS error of zero represents an unperturbed protein environment relative to the crystal structure. Well-preserved WS should be characterized by a low LAWS error. In contrast, a high LAWS error (> 3 Å^2^) represents a situation where the deviation of the protein structure is so considerable that the placement of the CWS becomes meaningless in a given frame.

The distributions of LAWS error for each of the three WS groups are presented as box plots in Fig. 6. These distributions are exponential, with high variance indicated by the long whiskers. The preserved CWS have a mean LAWS error (±*SD*) of 0.7 ± 2.2 Å^2^, the bulk-like WS have a mean value of 1.0 ± 2. 2 Å^2^, whereas the mean LAWS error for the obstructed WS is 1.8 ± 2.3 Å^2^. As expected, preserved CWS are characterized by relatively low LAWS errors (< 1 Å^2^) corresponding to more minor deviations of the local protein conformation from the crystal structure. In contrast, high LAWS errors are more commonly observed in obstructed and bulk-like WS, where the perturbations are more significant.

**Figure 6:**
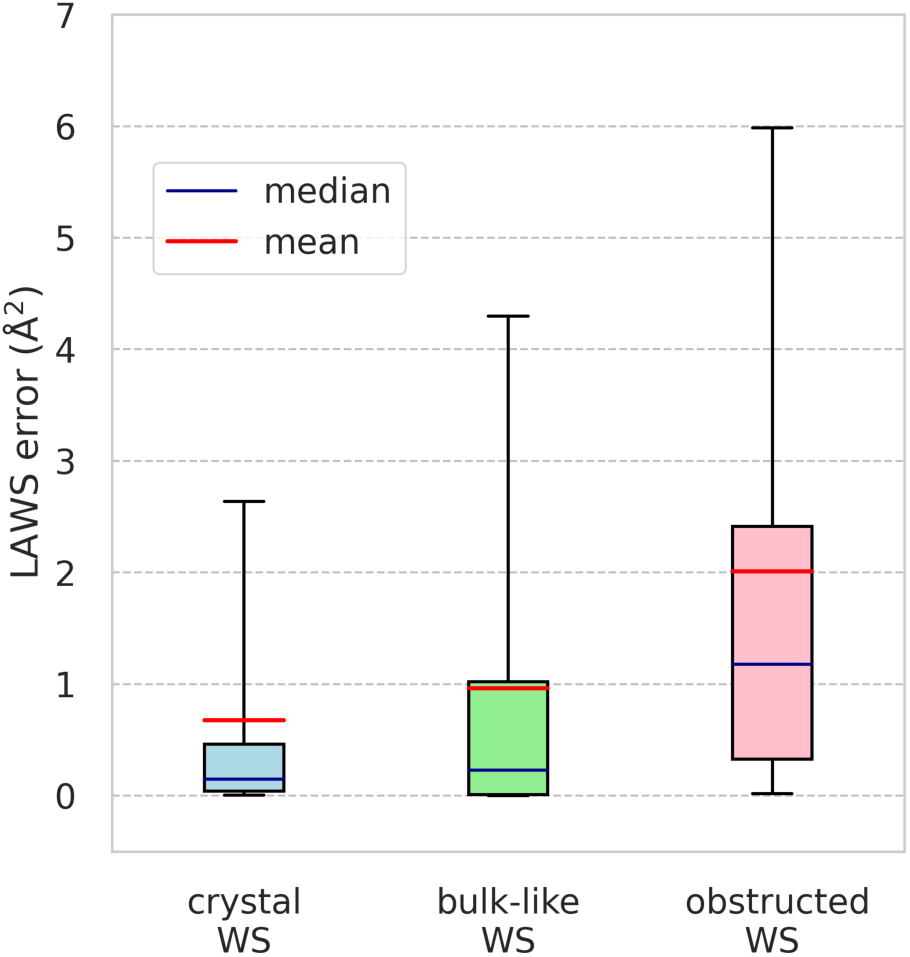
LAWS error for each category of water sites. The box plots show the distribution of LAWS error for crystal (blue), bulk-like (green) and obstructed (red) water sites. The LAWS error represents the minimized value of Eq.(2) when the CWS is optimally placed. In the plots, boxes represent the interquartile range (25th-75th percentile), while whiskers show 5th-95th percentile of the distribution.

We note that the LAWS error itself cannot be used as a criterion to determine if the CWS is preserved in a simulation. To illustrate this, we remove the frames for which the LAWS error exceeds 1 Å^2^ for each water site. By only analyzing the frames where CWS are well-coordinated (with a low LAWS error), we would expect perfect recall. Nevertheless, the observed recall with excluded frames is only slightly improved (79% as opposed to 76% for the entire trajectory). Therefore, low LAWS error is not a sufficient condition for CWS recall.

### Recall of CWS depends on the experimental B-factors

In diffraction experiments, B-factors are a measure of the spread of the electron density caused by atomic fluctuations in the crystal lattice. The experimental CWS (water oxygen atoms) have associated B-factors, which represent uncertainty in their positions. For this reason, we hypothesize that the experimental B-factor of a CWS should be related to the probability of preserving that CWS in the simulation.

To check whether this is the case, we analyzed CWS recall according to B-factor (Table 2). Using LAWS, 100% recall of CWS was obtained in the lowest B-factor range (< 20 Å^2^). For comparison, CWS in the middle range of 20-50 A have a recall of 74-81%, while CWS with high (> 50 Å^2^) B-factors are poorly preserved (0 to 67% recall). Additionally, the LAWS error correlates with B-factor, suggesting that the local protein environment surrounding the low B-factor CWS is less structurally perturbed than for the higher B-factor CWS.

**Table 2:** Recall of CWS in the simulation according to the experimental B-factor using LAWS and global alignment. The 94 CWS were grouped according to B-factor in bins of 10 Å^2^.

| B-factor range ($\text{\AA}^2$) | 10–20 | 20–30 | 30–40 | 40–50 | 50–60 | 60–70 |
| --- | --- | --- | --- | --- | --- | --- |
| # of CWS in range | 12 | 20 | 26 | 23 | 9 | 4 |
| Mean LAWS Error ($\text{\AA}^2$) | 0.23 | 0.85 | 0.88 | 1.04 | 0.50 | 2.45 |
| Preserved CWS using LAWS (%) | 100 | 75 | 81 | 74 | 67 | 0 |
| Preserved CWS using Global Alignment (%) | 92 | 65 | 65 | 61 | 78 | 50 |

An advantage of simulating the entire unit cell is the independent information provided by the four symmetric copies of each CWS (Fig. 3). Making use of this information, we computed the number of copies (from 0 to 4) that were classified as preserved for each of the 94 CWS (Fig. 7). More copies of the CWS with low B-factors are found to be preserved in the simulation. Similar to Table 2, this trend is robust regardless of whether LAWS or global alignment is used to track the water sites. Importantly, LAWS shows a higher consistency (across symmetric copies) than global alignment in the low B-factor range.

**Figure 7:**
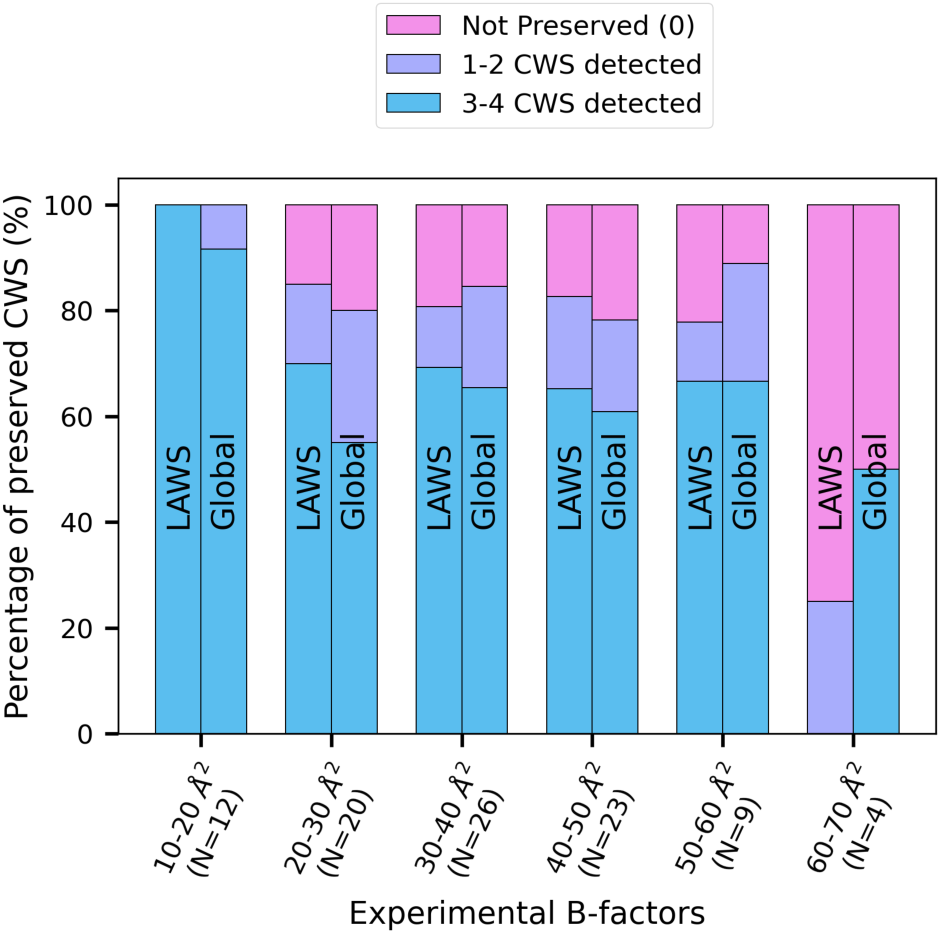
Experimental B-factors correlate with preserving CWS in the simulation. The bar chart shows the percentage of the CWS which are preserved in 0 (pink), 1-2 (purple), or 3-4 (blue) symmetric copies in the unit cell. Results are provided for both LAWS and the global alignment approach. The 94 CWS were grouped according to B-factor in bins of 10 Å^2^, with the number of CWS in each bin, N, provided.

Taken together, this analysis reveals that the CWS with lower B-factors, i.e. lower uncertainty, are more likely to be preserved in the simulation, as opposed to the CWS with higher B-factors, and higher associated uncertainty in their positions. This trend is consistent with the findings of Sun et al., who investigated the preservation of binding site waters in simulations.^23^ Similarly, we analyzed local 3D density peaks for preserved CWS, and a negative correlation between peak height and experimental B-factor is found (Fig. S4).

Interestingly, LAWS demonstrates a low recall of the highest uncertainty CWS, with B-factor > 60 Å^2^ (Table 2). These sites also have the highest LAWS error, which suggests that the surrounding protein structure is highly perturbed. It is unclear how many of these sites are expected to be captured by simulation, since B-factors of > 60 Å^2^ are close to the bulk threshold for this system. To understand these effects in more detail, we analyzed the locations of the lost CWS.

### Lost CWS are coordinated by flexible regions of the protein

In the analysis carried out so far, we have used LAWS to quantify CWS recall. Next, we examine how the preservation of a water site is affected by the surrounding protein structure. Since we expect the protein-protein interfaces to have a higher mobility than the regions within the protein, we compared the CWS coordinated by the corresponding regions. These are (i) intra-chain CWS, i.e. those making contacts with a single protein chain, and (ii) inter-chain CWS found at the interface between protein chains. Interestingly, the experimental B-factors of these two groups do not differ significantly, with *B*_*intra*_= 36 ±13 Å^2^ (*n* = 38) and *B* _*inter*_ = 36 ± 12 Å^2^ (*n* = 56). The CWS recall is also found to be comparable, with 76% preserved for the intra-chain CWS and 75% for the inter-chain CWS. These results suggest that the CWS at the protein interfaces are equally well preserved in the simulation as the CWS coordinated by a single protein chain.

Further analysis of the lost CWS identifies that the key similarity between them appears to be tied to their location relative to the protein. These CWS are mostly coordinated by the flexible structural elements, including the loops and the C-terminal tail (Table S2). These protein segments have a high root-mean-squared fluctuation (RMSF) in the simulation (Fig. 8), indicating that the loss of CWS can be attributed to the higher mobility of the protein regions coordinating these water sites. The relatively high LAWS error observed for bulk-like and obstructed WS suggest that the flexible regions of the protein also experience significant deviations relative to the crystal structure, leading to a loss of water coordination. To determine whether this observation is general or specific to the system studied here will require further investigation.

**Figure 8:**
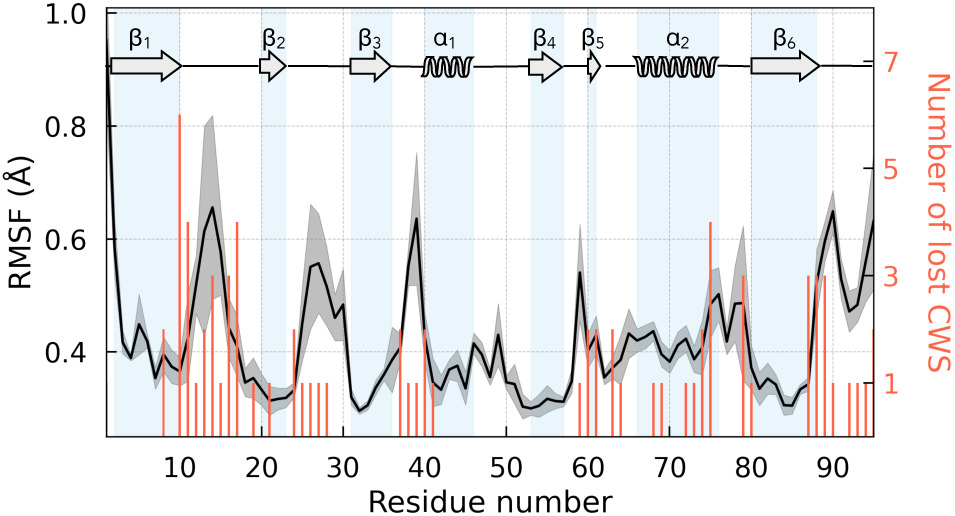
Lost CWS are coordinated by flexible regions of the protein. RMSF of C*α* atoms represented by the mean value (solid line) over *N* = 4 protein copies in the unit cell. The shaded envelope shows standard deviation. There are 23 lost CWS which are each coordinated by multiple protein residues. The number of the lost CWS coordinated by each residues is shown with red bars. This figure illustrates the data provided in Table S2.

## Conclusions

In this study, we discuss the limitations of existing global alignment approaches for analyzing crystallographic water in MD simulations. Namely, these methods are only suitable for cases when the protein conformation remains close to the crystal structure. To address this limitation of global alignment, we developed the LAWS method which reduces alignment error when protein regions experience significant deviations from the starting crystal structure.

To the best of our knowledge, the only study which implemented a similar local alignment approach was Caldararu et al. ^13^ We build upon their approach by using the radial density profile of water as the main criteria to determine the preserved CWS (instead of relying on data-driven clustering). In addition, our method utilizes a reference (bulk water sites) to decide whether or not a particular CWS is preserved. Importantly, LAWS takes into account the CWS coordinated by multiple symmetrically related protein monomers in the crystal lattice (Fig. 2A). This consideration is critical, since ∼60% of all CWS in the protein crystal simulated here are coordinated by protein atoms from multiple chains. The relative motion of adjacent chains, even if their structures remain stable, can perturb the coordination of CWS. Unique to our method, the LAWS error serves as a measure of protein structural perturbation around each water site; it quantifies how much the CWS deviates from its experimental position in each frame.

Applying the LAWS method to a 1 *µs* simulation of a protein crystal, we demonstrate that LAWS improves the recall of high-confidence (low B-factor) CWS and shows an increased density of water surrounding the CWS (Fig. 5, S4) compared to global alignment. We also use LAWS to compute CWS recall statistics. Previous studies reported the CWS recall as the fraction of water density peaks detected within 1.4 Å of their positions in the crystal structure. ^13–15,21^ They found that CWS recall varied from 40 to 100% depending on the system, simulation setup, and the method used for analysis. To place our results in the context of these studies, we provide analogous recall statistics computed using 3D density maps: 70% with LAWS vs 60% with global alignment (Table S1). This is broadly consistent with the earlier studies that did not use position restraints for a protein structure.

It is a question, what recall of CWS is realistic, given the challenges of accurately replicating the experiments. The lack of consensus between independent experiments in the positions of CWS^46–49^ suggests that complete recall of all CWS in simulations is most likely overly optimistic and should not be expected. The construction of the MD simulation system on its own has multiple challenges that can affect accuracy of modeling crystallographic water. For example, the resolution of the starting structure plays a crucial role in the quality of simulations.^50^ In addition, the correct assignment of protonation states^51^ and the inclusion of crowding agents in the crystal lattice^52^ can be challenging. Despite these difficulties, we observe 100% recall of the high confidence (low B-factor) CWS using LAWS in this study.

Since protein crystal simulations have been suggested as a benchmark system for force field development and testing,^26^ LAWS can be a valuable tool in comparing CWS recall statistics between force fields. Additionally, we envision that our method can be used to better characterize water networks in proteins, such as those found in the active site of enzymes^6^ or the conserved water networks in GPCRs.^53^ The multilateration approach described here could also be used to track the interactions of small molecules with flexible regions of proteins, such as binding pockets or allosteric sites.

## Acknowledgemen

This research was supported by a Connaught New Researcher Award to S.R., a Natural Science and Engineering Research Council of Canada (NSERC) Discovery Grant and by Compute Canada. Computations were performed on the Niagara supercomputer at the SciNet HPC Consortium. SciNet is funded by: the Canada Foundation for Innovation; the Government of Ontario; Ontario Research Fund - Research Excellence; and the University of Toronto.

## Supporting Information Available

1. Analysis of Bulk Water Sites: Figure S1
2. The LAWS pipeline
3. RMSD of the protein relative to the crystal structure: Figure S2
4. LAWS using local 3D density maps: Figure S3, Table S1, Figure S4
5. Locations of CWS that are not preserved: Table S2

Simulation data is available at https://doi.org/10.5281/zenodo.6478270

The implementation of the LAWS algorithm is available at https://github.com/rauscher-lab/LAWS

## Notes

https://doi.org/10.5281/zenodo.6478270

https://github.com/rauscher-lab/LAWS

### Competing Interest Statement

The authors have declared no competing interest.

## References

(1) Helms, V. Protein Dynamics Tightly Connected to the Dynamics of Surrounding and Internal Water Molecules. ChemPhysChem 2007, 8, 23–33, DOI: 10.1002/cphc.200600298.

(2) Bellissent-Funel, M.-C.; Hassanali, A.; Havenith, M.; Henchman, R.; Pohl, P.; Sterpone, F.; van der Spoel, D.; Xu, Y.; Garcia, A. E. Water Determines the Structure and Dynamics of Proteins. Chem. Rev. 2016, 116, 7673–7697, DOI: 10.1021/acs.chemrev.5b00664.

(3) Levy, Y.; Onuchic, J. N. Water Mediation in Protein Folding and Molecular Recognition. Annu. Rev. Biophys. Biomol. Struct. 2006, 35, 389–415, DOI: 10.1146/annurev.biophys.35.040405.102134.

(4) Schirò, G.; Fichou, Y.; Gallat, F.-X.; Wood, K.; Gabel, F.; Moulin, M.; Härtlein, M.; Heyden, M.; Colletier, J.-P.; Orecchini, A.; Paciaroni, A.; Wuttke, J.; Tobias, D. J.; Weik, M. Translational diffusion of hydration water correlates with functional motions in folded and intrinsically disordered proteins. Nat. Comm. 2015, 6, 6490, DOI: 10.1038/ncomms7490.

(5) Nakasako, M. Hydration Structures of Proteins; Springer Japan, 2021; DOI: 10.1007/978-4-431-56919-0.

(6) Kim, T. H.; Mehrabi, P.; Ren, Z.; Sljoka, A.; Ing, C.; Bezginov, A.; Ye, L.; Pom`es, R.; Prosser, R. S.; Pai, E. F. The role of dimer asymmetry and protomer dynamics in enzyme catalysis. Science 2017, 355, eaag2355, DOI: 10.1126/science.aag2355.

(7) Karplus, M.; Kuriyan, J. Molecular dynamics and protein function. Proc. Natl. Acad. Sci. U.S.A. 2005, 102, 6679–6685, DOI: 10.1073/pnas.0408930102.

(8) Berman, H. M.; Westbrook, J.; Feng, Z.; Gilliland, G.; Bhat, T. N.; Weissig, H.; Shindyalov, I. N.; Bourne, P. E. The Protein Data Bank. Nucleic Acids Res. 2000, 28, 235–242, DOI: 10.1093/nar/28.1.235.

(9) Cerutti, D. S.; Freddolino, P. L.; Duke, R. E.; Case, D. A. Simulations of a Protein Crystal with a High Resolution X-ray Structure: Evaluation of Force Fields and Water Models. J. Phys. Chem. B 2010, 114, 12811–12824, DOI: 10.1021/jp105813j.

(10) Janowski, P. A.; Cerutti, D. S.; Holton, J.; Case, D. A. Peptide Crystal Simulations Reveal Hidden Dynamics. J. Am. Chem. Soc. 2013, 135, 7938–7948, DOI: 10.1021/ja401382y.

(11) Cerutti, D. S.; Case, D. A. Molecular dynamics simulations of macromolecular crystals. Wiley Interdiscip. Rev. Comput. Mol. Sci. 2018, 9, e1402, DOI: 10.1002/wcms.1402.

(12) Case, D. A.; Cheatham, T. E.; Darden, T.; Gohlke, H.; Luo, R.; Merz, K. M.; Onufriev, A.; Simmerling, C.; Wang, B.; Woods, R. J. The Amber biomolecular simulation programs. J. Comput. Chem. 2005, 26, 1668–1688, DOI: 10.1002/jcc.20290.

(13) Caldararu, O.; Ignjatović, M. M.; Oksanen, E.; Ryde, U. Water structure in solution and crystal molecular dynamics simulations compared to protein crystal structures. RSC Adv. 2020, 10, 8435–8443, DOI: 10.1039/c9ra09601a.

(14) Wall, M. E.; Calabró, G.; Bayly, C. I.; Mobley, D. L.; Warren, G. L. Biomolecular Solvation Structure Revealed by Molecular Dynamics Simulations. J. Am. Chem. Soc. 2019, 141, 4711–4720, DOI: 10.1021/jacs.8b13613.

(15) Altan, I.; Fusco, D.; Afonine, P. V.; Charbonneau, P. Learning about Biomolecular Solvation from Water in Protein Crystals. J. Phys. Chem. B 2018, 122, 2475–2486, DOI: 10.1021/acs.jpcb.7b09898.

(16) Copie, G.; Cleri, F.; Blossey, R.; Lensink, M. F. On the ability of molecular dynamics simulation and continuum electrostatics to treat interfacial water molecules in proteinprotein complexes. Sci. Rep. 2016, 6, 38259, DOI: 10.1038/srep38259.

(17) Henchman, R. H.; McCammon, J. A. Extracting hydration sites around proteins from explicit water simulations. J. Comput. Chem. 2002, 23, 861–869, DOI: 10.1002/jcc.10074.

(18) Higo, J.; Nakasako, M. Hydration structure of human lysozyme investigated by molecular dynamics simulation and cryogenic X-ray crystal structure analyses: On the correlation between crystal water sites, solvent density, and solvent dipole. J. Comput. Chem. 2002, 23, 1323–1336, DOI: 10.1002/jcc.10100.

(19) Feig, M.; Pettitt, B. M. Crystallographic water sites from a theoretical perspective. Structure 1998, 6, 1351–1354, DOI: 10.1016/s0969-2126(98)00135-x.

(20) Steinbach, P. J.; Brooks, B. R. Protein hydration elucidated by molecular dynamics simulation. Proc. Natl. Acad. Sci. U.S.A. 1993, 90, 9135–9139, DOI: 10.1073/pnas.90.19.9135.

(21) Rudling, A.; Orro, A.; Carlsson, J. Prediction of Ordered Water Molecules in Protein Binding Sites from Molecular Dynamics Simulations: The Impact of Ligand Binding on Hydration Networks. J. Chem. Inf. Model. 2018, 58, 350–361, DOI: 10.1021/acs.jcim.7b00520.

(22) Young, T.; Abel, R.; Kim, B.; Berne, B. J.; Friesner, R. A. Motifs for molecular recognition exploiting hydrophobic enclosure in protein-ligand binding. Proc. Natl. Acad. Sci. U.S.A. 2007, 104, 808–813, DOI: 10.1073/pnas.0610202104.

(23) Sun, H.; Zhao, L.; Peng, S.; Huang, N. Incorporating replacement free energy of binding-site waters in molecular docking. Proteins 2014, 82, 1765–1776, DOI: 10.1002/prot.24530.

(24) van Gunsteren, W. F.; Berendsen, H. J.; Hermans, J.; Hol, W. G.; Postma, J. P. Computer simulation of the dynamics of hydrated protein crystals and its comparison with x-ray data. Proc. Natl. Acad. Sci. U.S.A. 1983, 80, 4315–4319, DOI: 10.1073/pnas.80.14.4315.

(25) Hekstra, D. R.; White, K. I.; Socolich, M. A.; Henning, R. W.; Šrajer, V.; Ranganathan, R. Electric-field-stimulated protein mechanics. Nature 2016, 540, 400–405, DOI: 10.1038/nature20571.

(26) Janowski, P. A.; Liu, C.; Deckman, J.; Case, D. A. Molecular dynamics simulation of triclinic lysozyme in a crystal lattice. Protein Sci. 2016, 25, 87–102, DOI: 10.1002/pro.2713.

(27) Rodriguez, M. Z.; Comin, C. H.; Casanova, D.; Bruno, O. M.; Amancio, D. R.; Costa, L. d. F.; Rodrigues, F. A. Clustering algorithms: A comparative approach. PLoS One 2019, 14, e0210236, DOI: 10.1371/journal.pone.0210236.

(28) Jaulin, L. Instantaneous Localization. In Mobile Robotics; Jaulin, L.; Ed., Elsevier, 2015; pp 171–196, DOI: 10.1016/B978-1-78548-048-5.50005-X.

(29) Moré, J. J. The Levenberg-Marquardt algorithm: Implementation and theory. In Lecture Notes in Mathematics; Springer Berlin Heidelberg, 1978; pp 105–116, DOI: 10.1007/bfb0067700.

(30) Briones, R.; Blau, C.; Kutzner, C.; de Groot, B. L.; Aponte-Santamaria, C. GROmaρs: A GROMACS-Based Toolset to Analyze Density Maps Derived from Molecular Dynamics Simulations. Biophys. J. 2019, 116, 4–11, DOI: 10.1016/j.bpj.2018.11.3126.

(31) Chen, X.; Weber, I.; Harrison, R. W. Hydration Water and Bulk Water in Proteins Have Distinct Properties in Radial Distributions Calculated from 105 Atomic Resolution Crystal Structures. J. Phys. Chem. 2008, 112, 12073–12080, DOI: 10.1021/jp802795a.

(32) Ebbinghaus, S.; Kim, S. J.; Heyden, M.; Yu, X.; Heugen, U.; Gruebele, M.; Leitner, D. M.; Havenith, M. An extended dynamical hydration shell around proteins. Proc. Natl. Acad. Sci. U.S.A. 2007, 104, 20749–20752, DOI: 10.1073/pnas.0709207104.

(33) Huang, Y.; Zhang, X.; Ma, Z.; Li, W.; Zhou, Y.; Zhou, J.; Zheng, W.; Sun, C. Q. Size; separation, structural order and mass density of molecules packing in water and ice. Sci. Rep. 2013, 3, 3005, DOI: 10.1038/srep03005.

(34) Bergmann, U.; Cicco, A. D.; Wernet, P.; Principi, E.; Glatzel, P.; Nilsson, A. Nearestneighbor oxygen distances in liquid water and ice observed by x-ray Raman based extended x-ray absorption fine structure. J. Chem. Phys. 2007, 127, 174504, DOI: 10.1063/1.2784123.

(35) Michaud-Agrawal, N.; Denning, E. J.; Woolf, T. B.; Beckstein, O. MDAnalysis: A toolkit for the analysis of molecular dynamics simulations. J. Comput. Chem. 2011, 32, 2319–2327, DOI: 10.1002/jcc.21787.

(36) Jo, S.; Kim, T.; Iyer, V. G.; Im, W. CHARMM-GUI: A web-based graphical user interface for CHARMM. J. Comput. Chem. 2008, 29, 1859–1865, DOI: 10.1002/jcc.20945.

(37) Abraham, M. J.; Murtola, T.; Schulz, R.; Pall, S.; Smith, J. C.; Hess, B.; Lindahl, E. GROMACS: High performance molecular simulations through multi-level parallelism from laptops to supercomputers. SoftwareX 2015, 1–2, 19–25, DOI: 10.1016/j.softx.2015.06.001.

(38) Huang, J.; Rauscher, S.; Nawrocki, G.; Ran, T.; Feig, M.; de Groot, B. L.; Grubmüller, H.; MacKerell, A. D. CHARMM36m: an improved force field for folded and intrinsically disordered proteins. Nat. Methods 2017, 14, 71–73, DOI: 10.1038/nmeth.4067.

(39) MacKerell, A. D.; Bashford, D.; Bellott, M.; Dunbrack, R. L.; Evanseck, J. D.; Field, M. J.; Fischer, S.; Gao, J.; Guo, H.; Ha, S.; Joseph-McCarthy, D.; Kuchnir, L.; Kuczera, K.; Lau, F. T. K.; Mattos, C.; Michnick, S.; Ngo, T.; Nguyen, D. T.; Prodhom, B.; Reiher, W. E.; Roux, B.; Schlenkrich, M.; Smith, J. C.; Stote, R.; Straub, J.; Watanabe, M.; Wiórkiewicz-Kuczera, J.; Yin, D.; Karplus, M. All-Atom Empirical Potential for Molecular Modeling and Dynamics Studies of Proteins. J. Phys. Chem. B 1998, 102, 3586–3616, DOI: 10.1021/jp973084f.

(40) Hess, B.; Bekker, H.; Berendsen, H. J. C.; Fraaije, J. G. E. M. LINCS: A linear constraint solver for molecular simulations. J. Comput. Chem. 1997, 18, 1463–1472, DOI: 10.1002/(sici)1096-987x(199709)18:12<1463::aid-jcc4>3.0.co;2-h.

(41) Essmann, U.; Perera, L.; Berkowitz, M. L.; Darden, T.; Lee, H.; Pedersen, L. G. A smooth particle mesh Ewald method. J. Chem. Phys. 1995, 103, 8577–8593, DOI: 10.1063/1.470117.

(42) Taulier, N.; Chalikian, T. V. Compressibility of protein transitions. Biochim. Biophys. Acta, Protein Struct. Mol. Enzymol. 2002, 1595, 48–70, DOI: 10.1016/s0167-4838(01)00334-x.

(43) Bussi, G.; Donadio, D.; Parrinello, M. Canonical sampling through velocity rescaling. J. Chem. Phys. 2007, 126, 014101, DOI: 10.1063/1.2408420.

(44) Berendsen, H. J. C.; Postma, J. P. M.; van Gunsteren, W. F.; DiNola, A.; Haak, J. R. Molecular dynamics with coupling to an external bath. J. Chem. Phys. 1984, 81, 3684–3690, DOI: 10.1063/1.448118.

(45) Parrinello, M.; Rahman, A. Polymorphic transitions in single crystals: A new molecular dynamics method. J. Appl. Phys. 1981, 52, 7182–7190, DOI: 10.1063/1.328693.

(46) Levitt, M.; Park, B. H. Water: now you see it, now you don’t. Structure 1993, 1, 223–226, DOI: 10.1016/0969-2126(93)90011-5.

(47) Otting, G.; Liepinsh, E.; Wüthrich, K. Protein Hydration in Aqueous Solution. In NMR in Structural Biology ; World Scientific, 1995; pp 632–638, DOI: 10.1142/9789812795830_0057.

(48) Zhang, X.-J.; Matthews, B. Conservation of solvent-binding sites in 10 crystal forms of T4 lysozyme. Protein Sci. 1994, 3, 1031–1039, DOI: 10.1002/pro.5560030705.

(49) Sanschagrin, P. C.; Kuhn, L. A. Cluster analysis of consensus water sites in thrombin and trypsin shows conservation between serine proteases and contributions to ligand specificity. Protein Sci. 1998, 7, 2054–2064, DOI: 10.1002/pro.5560071002.

(50) Garcia-Viloca, M.; Poulsen, T. D.; Truhlar, D. G.; Gao, J. Sensitivity of molecular dynamics simulations to the choice of the X-ray structure used to model an enzymatic reaction. Protein Sci. 2004, 13, 2341–2354, DOI: 10.1110/ps.03504104.

(51) Socher, E.; Sticht, H. Mimicking titration experiments with MD simulations: A protocol for the investigation of pH-dependent effects on proteins. Sci. Rep. 2016, 6, 22523, DOI: 10.1038/srep22523.

(52) Cerutti, D. S.; Le Trong, I.; Stenkamp, R. E.; Lybrand, T. P. Simulations of a protein crystal: explicit treatment of crystallization conditions links theory and experiment in the streptavidin-biotin complex. Biochemistry 2008, 47, 12065–77, DOI: 10.1021/bi800894u.

(53) Venkatakrishnan, A. J.; Ma, A. K.; Fonseca, R.; Latorraca, N. R.; Kelly, B.; Betz, R. M.; Asawa, C.; Kobilka, B. K.; Dror, R. O. Diverse GPCRs exhibit conserved water networks for stabilization and activation. Proc. Natl. Acad. Sci. U. S. A. 2019, 116, 3288–3293, DOI: 10.1073/pnas.1809251116.

