## Supporting Information for "LAWS: Local Alignment for Water Sites — tracking ordered water in simulations"

### Contents

**1 Analysis of Bulk Water Sites** . . . **S-3**

**2 The LAWS pipeline** . . . **S-5**

**3 RMSD of the protein relative to the crystal structure** . . . **S-7**

**4 LAWS using local 3D density maps** . . . **S-8**

**5 Locations of CWS that are not preserved** . . . **S-11**

**References** . . . **S-12**

### List of Figures

S1 Comparison of bulk water sites from the protein crystal and the water box simulations . . . S-3

S2 Protein RMSD relative to the crystal structure . . . S-7

S3 A difference density map generated by offset vectors . . . S-8

S4 Density peaks of CWS are negatively correlated with experimental B-factors S-10

### List of Tables

S1 Recall of CWS determined from density maps . . . S-9

S2 The protein regions associated with the lost CWS in the simulation . . . S-11

### 1 Analysis of Bulk Water Sites

In the LAWS algorithm, we track the positions of water sites and compute the local density of water in the vicinity of these positions. The local and bulk water densities are then compared to identify if a water site has a high enough water occupancy to be classified as crystallographic. Bulk water sites are positions within the crystal lattice that are sufficiently far from the protein that interactions with the protein do not significantly influence the water structure. Waters located (at least) 6 Å away from the protein surface have been shown experimentally to display bulk-like behaviour (in terms of the lifetime of hydrogen bonds).[S1] Hence, we randomly chose $N = 120$ locations at least 6 Å from the protein in the simulation box of a crystal unit cell to represent bulk water sites. We then found the distribution of the distances to the nearest neighbour water $P(r)$ for all bulk water sites combined (Fig. S1A).

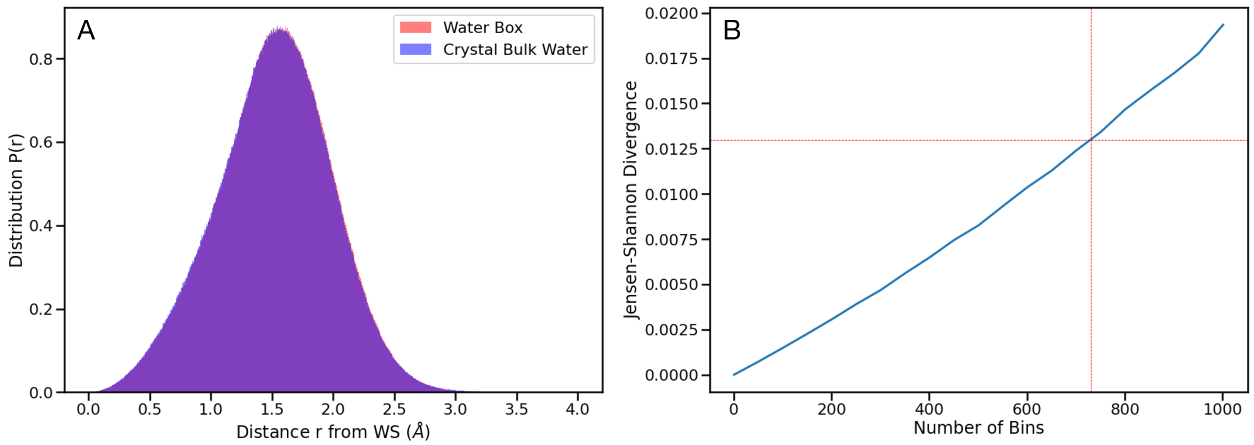


Figure S1: **Comparison of bulk water sites from the protein crystal and the water box simulations**. **(A)** Two (overlapped) histograms $P(r)$ were sampled from $N = 120$ bulk water sites in a water box simulation (red) and from the PDZ domain crystal simulation (blue). Here, $r$ represents the distance to the nearest-neighbour water molecule. **(B)** Jensen-Shannon divergence (JS) of these two histograms as a function of the number of bins chosen. The red dashed lines indicate the number of bins chosen (Freedman-Diaconis rule) and the corresponding JS score.

In order to check if the waters around these bulk water locations were indeed unaffected by the protein, we completed a similar analysis by sampling $N = 120$ random locations from an MD simulation of a box of water (simulation details provided below). As there is no protein in this system, all water molecules should have bulk-like behaviour. Fig. S1A shows the two $P(r)$ distributions for bulk-like sites from the crystal simulation and the simulation of the box of water. We compute the Jensen–Shannon (JS) divergence to quantify the difference between these two distributions. The JS divergence of these two distributions was 0.013, using the Freedman-Diaconis rule to assign the number of bins (Fig. S1B). Therefore, as the small JS divergence indicates the similarity of the distributions, our choice of bulk water sites is reasonable.

**Details of the simulation of water.** We constructed the simulation system with 8560 water molecules in a triclinic box of the same dimensions as the protein unit cell using the GROMACS utility *solvate*.[S2] The CHARMM-modified TIP3P water model[S3] was used for the simulation. The same equilibration steps were performed as for the protein crystal simulation, described in the Methods section of the main text. We performed one 500 ns production run of the system at 298 $K$ with isotropic Parrinello-Rahman pressure coupling.

#### 2 The LAWS pipeline

1. For each crystallographic water in the PDB:

   - Choose $n$ nearest-neighbour protein heavy atoms (within 4.5 Å of the crystallographic water).
   - Compute $\hat{d}_i$, the distances from each of the $n$ heavy atoms to the crystallographic water.
   - Compute weights $w_i^2 = \frac{(1/\hat{d}_i)^2}{\sum_{i=1}^{n}(1/\hat{d}_i)^2}$.

2. Sample $N = 120$ random bulk water sites in the unit cell, which are $\geq 6$ Å away from the protein.

3. At every simulation frame, collect statistics:

   (a) For each bulk water site:

   - Calculate the offset vector, $\vec{r}$, and the distance, $r = |\vec{r}|$, from the bulk WS to nearest water oxygen.

   (b) For each CWS:

   - Extract positions, $(x_i, y_i, z_i)$, for each of the $n$ heavy atoms.
   - Compute the new CWS position, $(x, y, z)$, by minimizing

   $$\mathrm{LAWS}(x, y, z) = \sum_{i=1}^{n} w_i^2 (\sqrt{(x - x_i)^2 + (y - y_i)^2 + (z - z_i)^2} - \hat{d}_i)^2.$$

   This minimum value is the LAWS error.

   - Calculate the offset vector, $\vec{r}$, and the distance, $r = |\vec{r}|$, from the CWS position to the nearest water oxygen.

4. For the calculated quantities ($\vec{r}$, $r$) corresponding to each CWS:

    - Compute the distance distribution, $P(r)$, and RDF, $g(r) = P(r)/r^2$.
    - Calculate the integral of $\int_0^{d_{OO}} g(r)dr$ and compare to the integral for bulk WS.
    - Classify the water site as a crystal, bulk-like, or obstructed WS (based on the integral value).

The implementation of the LAWS algorithm is publicly available: https://github.com/rauscher-lab/LAWS

### 3 RMSD of the protein relative to the crystal structure

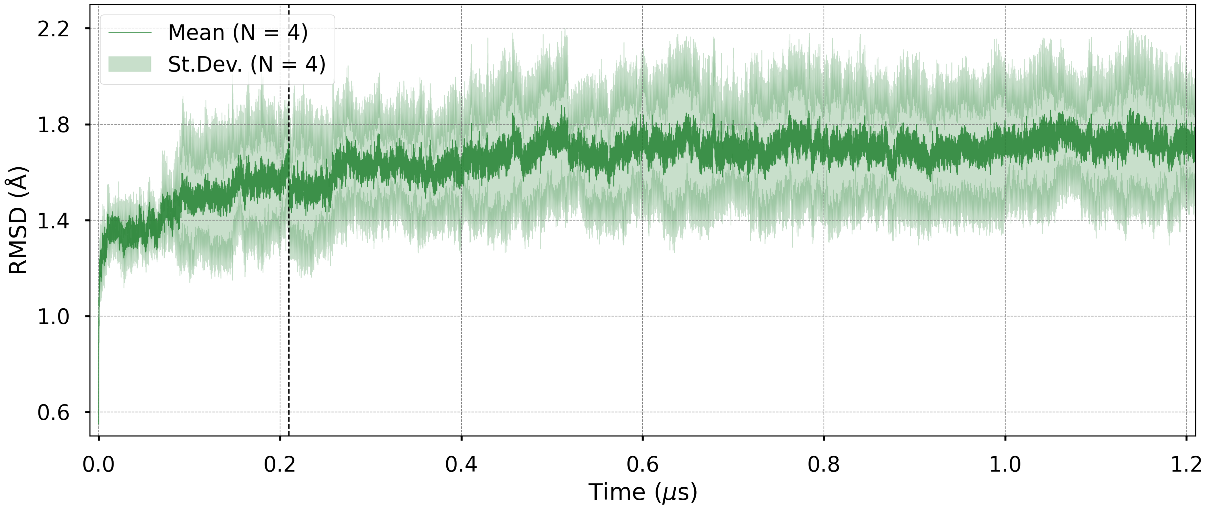


Figure S2: **Protein heavy-atom RMSD relative to the crystal structure as a function of time.** The mean RMSD (solid green line) over $N = 4$ chains and the standard deviation (shaded green envelope) is shown. The initial portion of the simulation was excluded from our analysis (black dashed line).

### 4 LAWS using local 3D density maps

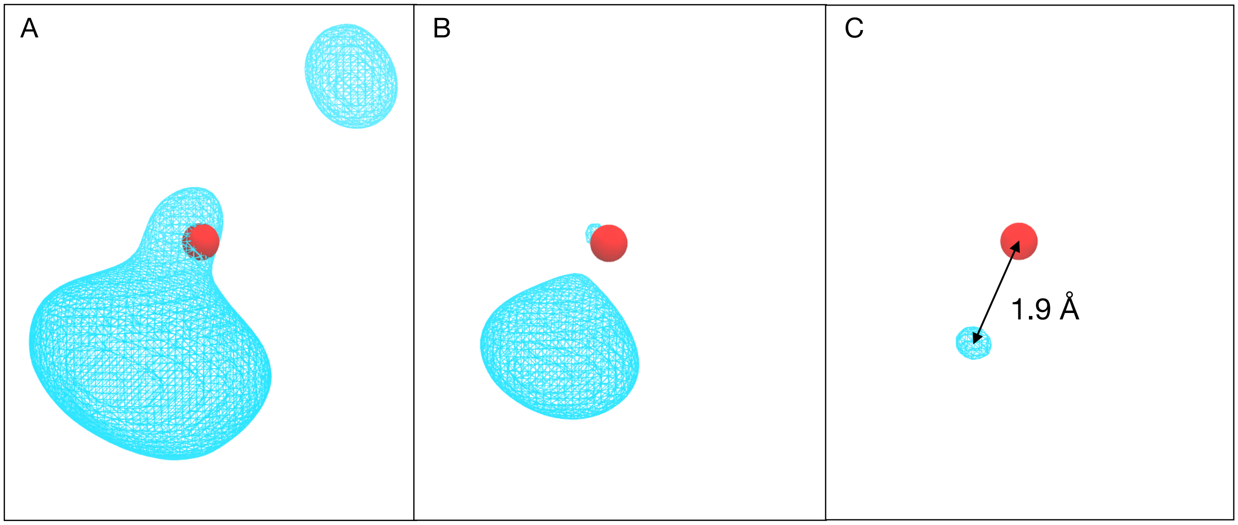


Figure S3: **A difference density map of the nearest neighbour water molecules relative to the bulk density**. Shown here is water site 49 (red sphere) with the density found using the LAWS method at varying thresholds. **(A)** A density isosurface of water at a threshold of bulk water. **(B)** A density isosurface representing the regions twice as dense as bulk. **(C)** The maximum peak, with a 6-fold greater density than bulk, located 1.9 Å from the water site.

The LAWS method provides a radial density profile of the nearest neighbour waters around a water site (Fig. 2 of the main text). However, it can be easily adapted to analyze 3D density maps around water sites tracked using LAWS. Offset vectors $\vec{r}$ around each water site contain $(x, y, z)$ coordinates of the closest water molecule at each simulation frame. First, we used the offset vectors from each frame to create an XTC trajectory for the water oxygens using MDAnalysis.[S4] Second, we used this trajectory as an input for the *maptide* command in the GROma$\rho$s software package[S5] to build a density map. Third, similar to the RDF analysis described in the main text, we compared the resulting density map to the reference density map obtained from combining the trajectory of the bulk water sites. We then applied the *mapdiff* utility in GROma$\rho$s to find a difference map. Finally, we identified the maximum positive peak (above the bulk water threshold) in a difference map that corresponds to the CWS (Fig. S3). The distance from the peak to the water site (i.e.

origin) was then measured. If no peaks are identified above the bulk water threshold, the water site is classified as bulk-like. The results of this analysis are presented in Table S1.

Table S1: **Recall of CWS in the 1-$\mu s$ MD simulation of the PDZ domain crystal determined from density maps using LAWS and global alignment methods.** Here, we present the number of CWS that have the highest density peak (above the bulk water threshold) within 0.5, 1.0, 1.4, 2.0, 2.5 and 3.0 Å (with the percentage of the total number $N = 94$ shown in parentheses). The number of bulk-like water sites is provided in the last column.

| Dist. to peak ($\mathring{A}$) | 0.5 | 1.0 | **1.4** | 2.0 | 2.5 | 3.0 | Bulk-like |
|---|---|---|---|---|---|---|---|
| LAWS | 29(31%) | 54(58%) | **66(70%)** | 74(79%) | 80(85%) | 83(88%) | 11(12%) |
| Global Alignment | 21(22%) | 44(47%) | **56(60%)** | 62(67%) | 66(70%) | 68(72%) | 23(24%) |

The recall of CWS at 1.4 Å found both by LAWS and global alignment (70% and 60%, respectively) is similar to the recall obtained using the radial distribution function approach (76% and 68%, respectively, reported in Table 1 in the main text). The number of bulk-like water sites found using density maps is comparable for LAWS (12% vs. 11% obtained using the RDF approach), while it is considerably higher for global alignment (24% vs 9% obtained using the RDF approach). It is important to note that when using the 3D-density rather than the RDF profile, none of the water sites are classified as obstructed.

We also examined if the height of the density peaks correlate with experimental isotropic B-factors for preserved CWS. Consistent with the results shown in Fig. 7, a negative correlation (Pearson correlation coefficient of $-0.7$) is observed for both the LAWS and global alignment density peaks. However, the density peaks obtained using LAWS are higher on average (Fig. S4) than the peaks obtained using global alignment, similar to the results shown in Fig. 5 in the main text.

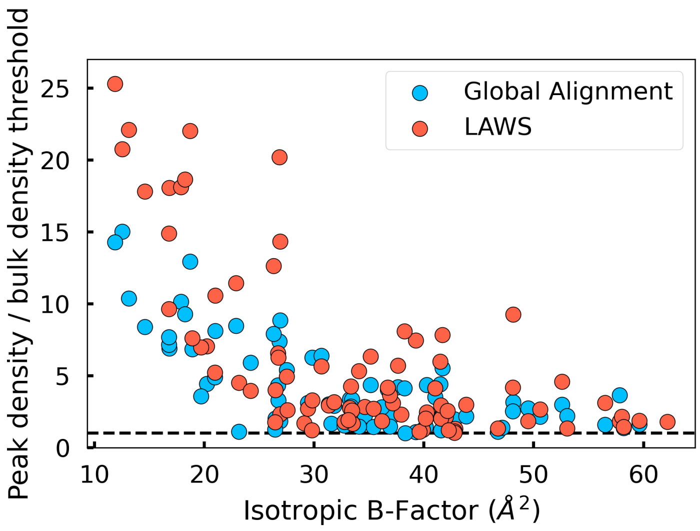


Figure S4: **Density peaks of CWS are negatively correlated with experimental B-factors**. LAWS detects higher density peaks on average when compared to global alignment. The peaks on the y-axis are normalized using the bulk density threshold.

### 5 Locations of CWS that are not preserved

Table S2: **The protein regions associated with the lost CWS in the simulation**. CWS are numerated consecutively as found in crystal structure (PDB 5E11). There are 9 intra-chain and 14 inter-chain CWS that were lost in the simulation. The intra-chain CWS are those coordinated by the regions found within the same protein chain. The inter-chain CWS are those coordinated by multiple protein chains. Note that the active site of the PDZ domain is a pocket between the $\alpha_2$-helix and the $\beta_2$-strand. For each CWS, at least one of the coordinating regions is a flexible loop/turn or the C-terminus, which also has a high RMSF. Exceptions to this rule include CWS with ids 91, 77, and 62, which are located near the $\alpha_2$ helix and $\beta_1$ strand.

| id | Type | Location |
|---|---|---|
| 0 | intra-chain | between $\beta_1$ and $\beta_6$ strands (close to $\beta_1$-$\beta_2$ loop) |
| 76 | intra-chain | near $\beta_5$-$\alpha_2$ loop |
| 79 | intra-chain | between $\beta_2$-$\beta_3$ loop and C-term. |
| 90 | intra-chain | between side-chain of $\beta_1$ strand and $\alpha_2$-$\beta_6$ loop |
| 29 | intra-chain | near $\beta_5$-$\alpha_2$ loop |
| 30 | intra-chain | between $\beta_1$-$\beta_2$ loop and $\alpha_2$ helix |
| 49 and 74 | intra-chain | between $\alpha_2$-$\beta_6$ and $\beta_1$-$\beta_2$ loops |
| 91 | intra-chain | between side chains of Gln72 and Ile73 ($\alpha_2$) |
| 1 and 5 | inter-chain | between $\beta_1$-$\beta_2$ loop and $\beta_3$-$\alpha_1$ loop |
| 7 | inter-chain | between $\alpha_2$ helix and C-term |
| 9 | inter-chain | between $\beta_2$-$\beta_3$ loop and C-term. |
| 17 | inter-chain | between $\beta_1$-$\beta_2$ loop and C-term. (in the active site) |
| 77 | inter-chain | between $\beta_1$ strand and $\beta_6$ strand and $\alpha_2$ helix |
| 93 | inter-chain | between $\beta_2$-$\beta_3$ loop and $\alpha_2$ helix |
| 8 | inter-chain | between $\beta_2$-$\beta_3$ loop and $\beta_6$ strand |
| 15 and 87 | inter-chain | between C-term. and $\alpha_2$ helix + $\beta_1$-$\beta_2$ loop |
| 86 | inter-chain | between $\beta_5$ strand and $\beta_1$-$\beta_2$ loop |
| 88 | inter-chain | between $\beta_2$-$\beta_3$ loop and $\beta_6$ strand |
| 25 | inter-chain | between $\beta_1$-$\beta_2$ loop and $\beta_4$-$\beta_5$ turn |
| 62 | inter-chain | between $\beta_1$ strand and $\beta_5$ strand |

### References


(S1) Ebbinghaus, S.; Kim, S. J.; Heyden, M.; Yu, X.; Heugen, U.; Gruebele, M.; Leitner, D. M.; Havenith, M. An extended dynamical hydration shell around proteins. *Proc. Natl. Acad. Sci. U.S.A.* **2007**, *104*, 20749–20752, DOI: 10.1073/pnas.0709207104.

(S2) Abraham, M. J.; Murtola, T.; Schulz, R.; Páll, S.; Smith, J. C.; Hess, B.; Lindahl, E. GROMACS: High performance molecular simulations through multi-level parallelism from laptops to supercomputers. *SoftwareX* **2015**, *1-2*, 19–25, DOI: 10.1016/j.softx.2015.06.001.

(S3) MacKerell, A. D.; Bashford, D.; Bellott, M.; Dunbrack, R. L.; Evanseck, J. D.; Field, M. J.; Fischer, S.; Gao, J.; Guo, H.; Ha, S.; Joseph-McCarthy, D.; Kuchnir, L.; Kuczera, K.; Lau, F. T. K.; Mattos, C.; Michnick, S.; Ngo, T.; Nguyen, D. T.; Prodhom, B.; Reiher, W. E.; Roux, B.; Schlenkrich, M.; Smith, J. C.; Stote, R.; Straub, J.; Watanabe, M.; Wiórkiewicz-Kuczera, J.; Yin, D.; Karplus, M. All-Atom Empirical Potential for Molecular Modeling and Dynamics Studies of Proteins. *J. Phys. Chem. B* **1998**, *102*, 3586–3616, DOI: 10.1021/jp973084f.

(S4) Michaud-Agrawal, N.; Denning, E. J.; Woolf, T. B.; Beckstein, O. MDAnalysis: A toolkit for the analysis of molecular dynamics simulations. *J. Comput. Chem.* **2011**, *32*, 2319–2327, DOI: 10.1002/jcc.21787.

(S5) Briones, R.; Blau, C.; Kutzner, C.; de Groot, B. L.; Aponte-Santamaria, C. GROma$\rho$s: A GROMACS-Based Toolset to Analyze Density Maps Derived from Molecular Dynamics Simulations. *Biophys. J.* **2019**, *116*, 4–11, DOI: 10.1016/j.bpj.2018.11.3126.
